# Predatory bacteria as members of human microbiomes and their impact on gut diversity and homeostasis

**DOI:** 10.64898/2026.07.03.736386

**Authors:** Johannes Zimmermann, Julia Johnke

## Abstract

**Background:** *Bdellovibrio* and like organisms (BALOs) are obligate bacterial predators that shape microbial communities by promoting species diversity, yet they have long been considered irrelevant to the human gut due to their presumed obligate aerobic lifestyle. Here, we challenge this view through a combined meta-analytic, experimental, and conceptual investigation of BALOs in human microbiomes.

**Results:** Reanalyzing 168,000 consistently processed samples from the Human Microbiome Compendium spanning 482 studies, we detected BALOs in more than 80 studies and across multiple body sites worldwide, with a gut prevalence of 2.4%, a finding confirmed by reanalysis of the PRIME database for 16S rRNA microbiome data. Strikingly, BALO presence was consistently associated with higher microbial alpha-diversity across body sites and disease contexts. Biopsy-derived samples showed a substantially higher prevalence than fecal samples, suggesting a mucosa-proximal niche. Our laboratory experiments showed that multiple *Bdellovibrio* strains can delay the loss of microbial diversity in vitro and remain active under gut-relevant conditions, including 37°C, pH 6.5, and in the presence of mucus. Genomic analyses further revealed terminal reductases, including nitrite reductases, in several BALO genomes, indicating the capacity for anaerobic or microaerobic respiration, consistent with persistence in mucosal microenvironments.

Notably, the metabolic and ecological profiles of cultured BALOs closely match those of facultative anaerobes, which constitute their preferred prey and are central drivers of dysbiosis in inflammatory bowel disease, diabetes, colorectal cancer, and chronic kidney disease.

**Conclusion:** Building on these findings, we propose a conceptual framework in which BALOs contribute to gut homeostasis by controlling the expansion of facultative anaerobes under inflammatory conditions, thereby facilitating the restoration of fermentative, butyrate-producing communities. Together, our results establish BALOs as consistent, functionally relevant members of the human microbiome and a promising natural candidate for therapeutic strategies targeting chronic gut disease.

## Background

The human gut microbiome is a complex and dynamic ecosystem in which microbial communities interact with one another and with the host. Most importantly, the microbiome has been reported to be involved in five out of eight hallmarks of health [1]. Thus, it is unsurprising that a microbial imbalance (called dysbiosis) is frequently observed in many diseases [2]. However, what constitutes a healthy versus a diseased microbiome remains unclear [3]. Microbiomes are individual and dynamic, making it difficult to define a single stable state associated with health or disease [4, 5].

Although some diseases have been linked to increased microbial diversity [6–8], the loss of diversity is generally considered a major factor contributing to dysbiosis and is therefore often used as an indicator distinguishing a “healthy” from a “diseased” microbiome [9–11]. Inflammatory bowel disease (IBD), for example, a chronic gut inflammation, alters the microbial composition [12], and studies have shown that the microbiomes of IBD patients are less diverse [13–15].

Interestingly, ecological principles can help explain the positive impact of diversity, since higher diversity has long been shown to enhance invasion resistance [16, 17] and ecosystem productivity [18, 19]. In general, microbiome redundancy has been described as a trait of human microbiomes [20], and increased diversity could enhance temporal stability as predicted by the insurance hypothesis [21]. In human microbiomes, the prominent loss of beneficial strains due to industrialization and modern lifestyles has been suggested as a contributor to many non-communicable diseases, in which a less diverse but well-adapted microbiome encounters a less-adapted, increasingly incompatible host [22–24]. Similarly, pathogens could be kept at a distance by more diverse microbial communities that prevent pathogenic colonization, for example, through competition for available nutrients [17]. In summary, although the link between microbiome diversity and health requires careful consideration [3], changes in diversity are affecting ecosystems and microbiomes in many ways.

Recent studies have highlighted BALOs obligate predatory bacteria that feed on Gram-negative bacteria as potential regulators of gut microbiome composition [25, 26]. BALOs prey either by invading the periplasmic space of their prey (i.e., periplasmic predation) or by vampirizing their prey while remaining outside the cell (i.e., epibiotic predation) [27]. Nowadays, they are collectively classified within the phylum Bdellovibrionota, which comprises BALOs from different genera that inhabit various environments, including soil, freshwater, and saltwater [28]. Due to their predatory nature, they have been proposed as “living antibiotic” [29] and have been used to treat specific bacterial infections (including [30–36]).

Notably, the presence of BALOs has been positively correlated with microbiome diversity across various hosts and environments, although not in the human gut [37, 38]. The putative “probiotic” effect is likely driven by top-down control within microbial communities, where predation reduces the abundance of dominant taxa, thereby freeing ecological niches and promoting diversity [39]. Alternatively, BALO predation might lead to changes in microbiome structure by enriching functionally important members, as demonstrated in the *C. elegans* microbiome, where BALO-driven community shifts increased vitamin B12 availability and significantly extended nematode lifespan [40].

An indication that a similar mechanism may operate in humans comes from a study reporting a positive association between BALOs and intestinal health. Specifically, reduced abundance or absence of BALOs in intestinal biopsy samples was associated with inflammatory bowel disease (IBD) [41]. However, despite this indirect evidence, empirical support for a positive effect of BALOs on human gut microbiome diversity and, ultimately, host health remains limited.

Given the described positive diversity effect and its potential links to health and ecosystem functioning, a key open question is whether BALO predation is driven by strain-level specificity, broad prey generalism, or selective targeting of ecologically or functionally defined bacterial groups. Usually, BALOs are described as exhibiting a broader prey range, in contrast to bacteriophages, which are usually highly host-specific [42] (but also see evidence for broader phage prey range and narrower BALO prey range [43, 44]). Interestingly, the commonly used type strain *Bdellovibrio bacteriovorus* HD100 can infect a wide range of Gram-negative bacteria, including facultative anaerobes such as *Escherichia coli, Salmonella, Pseudomonas,* and *Klebsiella* spp. [32, 35, 45], which are frequently associated with gut-related pathologies in humans [46].

Given this combination of ecological potential and clinical relevance, we focused our study on predators within the BALO group. Historically, BALOs have been rarely identified in human gut microbiome samples [41, 47], leading to the hypothesis that these predatory bacteria may be ill-suited to survive under the specific environmental stressors present in the gut, such as high temperatures (up to 37°C), low pH (down to 5.7), and primarily anoxic conditions, especially in the large intestine [48]. This is in line with the observation that higher abundances of *Bdellovibrio* were found within the duodenum and that the abundance decreases consistently towards the rectum [41]. Whether BALOs can survive and drive diversity under these conditions remains an open question.

Here, we aim to explore this conundrum by: (i) evaluating if BALO presence has a positive effect on microbial community diversity in a simple *in vitro* experiment, (ii) tracing BALO reads in human microbiomes, focusing on gut microbiomes, leveraging the newly available consistently processed microbial datasets such as the Human Microbiome Compendium (HMC) [49], (iii) identifying correlations between BALO presence, microbiome diversity, and health status for these subjects using the PRIME database with improved metadata [50], (iv) investigating the survival and activity of BALOs under gut-like conditions using laboratory experiments and genome-based predictions, and (v) we hypothesize that BALOs may inhabit the human gut microbiome and help restore dysbiotic community states by selectively targeting facultative anaerobes. Rethinking the role of BALOs in gut microbiomes is very promising, as it may provide new insights into the balance between microbial predation and the pathophysiology of gut diseases. If BALOs can survive under oxygen-limited conditions and prey on potentially harmful bacteria such as Proteobacteria, they could act as natural regulators of microbial populations, offering a novel avenue for therapeutic intervention in dysbiotic states.

## Material and Methods

### in vitro diversity experiment

#### BALO Cultivation

BALO isolates *Bdellovibrio tiberii* MYbb2 and *Bdellovibrio krueschi* MYbb4 [44] were cultivated following Jurkevitch et al. [51], with modifications. Briefly, 50 mL of 25 mM HEPES buffer (supplemented with 2 mM CaCl_2_, 3 mM MgCl_2_) was inoculated with either 1 mL of *E. coli* ML35 culture (MYbb2) or *Ochrobactrum vermis* MYb10 culture (MYbb4) at an OD_600_ of 10 and 200 μL of *Bdellovibrio* filtrate. The filtrate was obtained from a fresh BALO culture that had been passed through a 0.45 μm filter to remove residual prey cells. Cultures were incubated at 28°C with shaking for 24–48 hours. The resulting BALO culture, showing visible clearance, was again filtered through a 0.45 μm membrane to eliminate remaining prey cells. The final concentration was approximately 1 × 10⁵ cells/mL.

#### Bacterial Community (CeMbio) Cultivation

All CeMBio isolates were cultured for 24 hours as previously described [52, 53]. After centrifugation, bacterial pellets were resuspended in pre-chilled (4°C) HEPES buffer to a final concentration of approximately 1 × 10^7^ cells/mL. Cell concentrations for each isolate were determined individually by CFU counts. In brief, each isolate was grown overnight in LB medium, adjusted to OD_600_ = 5 using a photometer, then diluted 1:10 and serially diluted from 10^−1^ to 10^−8^. Triplicates of 10 μL per dilution were plated on LB agar and incubated at 28°C overnight. CFU counts were used to calculate the dilution factor required to achieve 1 × 10^7^ cells/mL for each isolate.

A mixture of all CeMBio isolates was then prepared by combining equal numbers of cells from each isolate. BALO cultures (prepared as above) were added to this community after being stored at 4°C for 2 days.

#### Experimental Procedure

The experiment ran for 12 days in cell culture flasks (Sarstedt, #83.3911.502) containing 20 mL of 1:10 diluted LB. To avoid nutrient depletion, 10% of the bacterial suspension was transferred into fresh 1:10 diluted LB medium every 12 hours. These transfer frequency and dilution ratio were selected to minimize bottleneck effects while ensuring continuous nutrient supply. Each treatment group was maintained in six biological replicates.

#### DNA Extraction and Sequencing

To analyze the effect of BALO exposure on microbial diversity over time, 1 mL of culture was collected at each sampling point. Bacterial pellets were processed for DNA extraction using a CTAB-based extraction protocol [54].

#### Amplicon Sequencing and Raw Data Processing

Partial 16S rRNA gene sequences were amplified from the V3-V4 hypervariable region using the primers 341F (5′-CCTACGGGNGGCWGCAG-3′) and 805R (5′-GACTACHVGGGTATCTAATCC-3′) [55, 56]. Sequencing was performed on an Illumina MiSeq platform using a paired-end approach. Raw sequencing reads were first processed to remove adapter and primer sequences using Cutadapt (v4.4) [57]. The resulting pre-trimmed FASTQ files were processed using the DADA2 pipeline (v1.28) [58] in R (v4.3) to infer Amplicon Sequence Variants (ASVs). Forward and reverse read quality profiles were inspected visually, and reads were filtered and trimmed using filterAndTrim() with the following parameters: forward reads truncated to 270 bp, reverse reads to 205 bp, maximum expected errors of 2 (maxEE = c(2, 2)), and a truncation quality threshold of 2 (truncQ = 2). Bases with ambiguous assignments were prohibited (maxN = 0), and PhiX reads were removed. Error rates were estimated separately for forward and reverse reads using learnErrors(). After sample inference with the core DADA2 algorithm, paired reads were merged using mergePairs(). Amplicons outside the expected length range of 402–428 bp were removed. Chimeric sequences were identified and removed using removeBimeraDenovo() via the consensus method.

#### Taxonomic Assignment and CeMbio Mapping

Taxonomic classification was performed using assignTaxonomy() against the SILVA reference database (v138.2, nr99), followed by species-level assignment using addSpecies(). To identify ASVs corresponding to the *C. elegans* microbiota (CeMbio12 and CeMbio43, [53, 59]), representative ASV sequences were queried against a custom nucleotide BLAST database (using makeblastdb, v2.16.0+) of CeMbio 16S rRNA sequences using BLASTn [60]. Matches were retained at a threshold of ≥99.5% sequence identity and ≥95% query coverage. Accordingly, multiple ASVs were allowed to map to the same CeMbio strain to account for potential sequencing errors or intra-strain evolutionary divergence. ASVs not matching known CeMbio sequences at this threshold were categorized as “other.”

#### Community Filtering and Decontamination

Downstream analysis was conducted using the phyloseq package (v1.48.0) [61]. Negative controls and mock communities were used to identify potential contaminants through the prevalence-based method in the decontam package (v1.22.0) [62]. ASVs identified as contaminants or consistently detected in negative controls were manually excluded. Following decontamination, control samples were removed, and samples with fewer than 1,000 total reads were discarded.

Mitochondrial and chloroplast sequences were excluded. To improve the robustness of richness estimates, low-abundance taxa were filtered: ASVs were retained only if they were present in more than three samples, had a total abundance >5 reads across the dataset, and reached a maximum abundance >2 reads in at least one sample.

#### Diversity and Statistical Analysis

Observed ASV richness was calculated using phyloseq [61]. To evaluate non-predatory community richness, ASVs taxonomically assigned to *Bdellovibrio*-and-like organisms (BALOs) were identified by pattern matching in the taxonomy table and subtracted from the total observed richness on a per-sample basis.

Statistical analysis was performed using Generalized Linear Mixed Models (GLMMs) implemented in the glmmTMBpackage (v1.1.11) [63]. Temporal changes in richness were modeled using the interaction between treatment (e.g., BALO presence/absence or specific BALO strain) and time (modeled as a categorical factor), with ‘biological replicate’ included as a random intercept. The appropriate error distribution (Gaussian or Conway-Maxwell-Poisson) was selected for each analysis based on residual diagnostics using the DHARMa package (v0.4.7) [64], ensuring no significant deviations in dispersion, outliers, or residual patterns.

Global effects were assessed using Type II Wald tests, and post-hoc pairwise comparisons between treatments at each timepoint were conducted using estimated marginal means (emmeans, v1.11.1) [65]. P-values were adjusted for multiple testing using the Benjamini–Hochberg (FDR) procedure. All visualizations were generated using ggplot2 (v3.5.2) [66], with error bars representing the standard error of the mean (SEM).

### database analyses

#### HMC

We utilized the Human Microbiome Compendium (HMC) [49] v1.1.0, released in September 2024 and based on SILVA v138.2 (https://doi.org/10.5281/zenodo.8186993), for our analysis. This resource comprises 168,000 consistently processed gut microbiome samples from 482 projects worldwide, spanning 68 countries. BALOs were defined as ASVs assigned to the genera *Bdellovibrio*, *Pseudobdellovibrio*, *Bacteriovorax*, *Peredibacter, Halobacteriovorax*, or *Pseudobacteriovorax* within the phylum *Bdellovibrionota* or the genus *Micavibrio* here still classified within the phylum *Pseudomonadota*. We considered only samples with 1000 < total_reads < 100000, resulting in 83 projects (i.e., studies) containing BALO reads (i.e., a project with at least one sample having > 0 BALO reads) with 909 samples having BALOs and 36,468 without BALOs.

#### PRIME

As validation, we investigated the Phenotypic Reference for Integrated Microbiome Enrichment (PRIME) [50] v1 released in July 2025 and based on SILVA v138.2 (https://zenodo.org/records/20200953) and repeated our analysis. This resource contains 53,449 consistently processed microbiome samples from 129 projects worldwide, spanning 85 countries. Compared to the HMC dataset, the sample-level metadata were manually curated and standardized across studies, and more than gut microbiomes were included. BALOs were defined as ASVs assigned to the genera *Bdellovibrio*, *Bacteriovorax*, *Peredibacter, Halobacteriovorax*, or *Pseudobacteriovorax* within the phylum *Bdellovibrionota* or the genus *Micavibrio,* here still classified within the phylum *Proteobacteria.* In contrast to the HMC analysis, *Pseudobdellovibrio was* not found within the PRIME taxonomy. We considered only samples with a quality tag of “Excellent” or “Good” in the metadata and identified 57 projects containing BALO reads (i.e., a project with at least one sample with> 0 BALO reads), resulting in 1809 samples with BALOs and 44542 samples without BALOs.

#### Diversity analysis

To account for differences in sequencing depth, which could directly influence BALO detection and diversity estimates, we compared median read counts per project. Distributions of read counts from BALO and no-BALO samples were similar (Fig. S1), indicating that additional reads were not responsible for detecting BALOs in the samples. Next, we calculated the mean and median alpha diversity (Chao, Shannon, Simpson) of each project’s BALO and non-BALO samples using the R package vegan v2.7.3 [67]. We also compared read counts across samples to assess the influence of sequencing depth. To compare the diversity of BALO and no BALO samples, we, whenever possible, compared BALO project medians using the Wilcoxon test for paired data to avoid the impact of larger studies and differences in read depth. In some cases, when the number of projects was low (e.g., for the diversity effect in body sites or disease types), we compared samples directly. In those cases, we compared differences in alpha diversity using an unpaired Wilcoxon test (P-values were BH-corrected) and, more importantly, the overall BALO effect using a linear mixed-effects model with *BALO* (true/false) as a fixed effect and *study number* and *read count* as random effects. In particular, the design formula for the model was *lmer(chao ∼ balo2 + (1|project) + (1|total_reads)* and the lmerTest R package v3.2-1 was employed [68].

#### World map

We created a map of found BALO samples worldwide by using the *geo_loc_name* tag for each sample in the HMC dataset (https://zenodo.org/records/15122187, file: tags.tsv.gz) and the R package sf [69] v.1.1-1. The number of samples found per country was grouped into ranges and plotted using ggplot2 v.4.0.3.

#### Oxygen preference trait prediction

We first downloaded all ASV sequences of the HMC dataset from the HMC Zenodo reference (https://zenodo.org/records/15122187, file: *obs_md.txt.gz*). The oxygen preference of all ASVs of the HMC dataset was then predicted with AmpliconTraits [70] using usearch v11.0.667 (parameter: *-sinaps -strand plus*) and the AmpliconTraits database *metabolism.fasta* (downloaded from https://github.com/jdonhauser/ampliconTraits on 2024/01/19). That resulted in a file *hmc-metabolism_amplicontraits.txt.gz,* which contains the oxygen preference for each ASV. Finally, we aggregated the relative abundances of all ASVs belonging to the same groups (anaerobic, facultative, microaerophilic, obligate_anaerobic, aerobic, obligate_aerobic) for each sample to obtain per sample-wise oxygen preferences. In addition, the facultative anaerobe trait of Table S2 was derived from the literature, mainly the Madin and BacDive databases [71, 72].

#### Network analysis

Networks were generated using all Human Microbiome Project data from the HMC. ASV count data were first converted to relative abundances to remove all ASVs with mean relative abundances below 0.001%. This ASV table was then split into samples containing BALO reads and those without. From both sets, ASVs with zero reads were removed. This resulted in two ASV tables: one with BALO-containing samples (1110 samples with 722 ASVs) and one without BALO-containing samples (55287 samples with 725 ASVs). Microbial co-occurrence correlations were estimated using FastSpar [73], which accounts for compositionality in microbiome data. Median correlation and covariance matrices were computed from the respective ASV table, and 1,000 bootstrap resamples were generated to assess variability. Empirical p-values were calculated from the bootstrap correlations using 1,000 permutations, and correlations with p < 0.05 were considered statistically significant. We further filtered edges between ASVs (i.e., a significant positive or negative correlation) based on different correlation thresholds and visualized networks using the R package igraph [74] (version 2.1.4).

### survival under gut-like conditions

#### Predation under gut-like pH and temperature

Predation by *Bdellovibrio* on prey *E. coli* ML35 was tested under different conditions using the predators *B. bacteriovorus* HD100, *B. tomkyle* MYbb1, *B. tiberii* MYbb2, *B. kumpostii* MYbb5, and *B. bagaluti* MYbb7. *Bdellovibrio* strains MYbb1, MYbb2, MYbb5, and MYbb7 were isolated from the natural environment of *Caenorhabditis elegans*. Furthermore, their 16S rRNA gene sequences are frequently detected in *C. elegans* microbiome datasets, indicating that these strains are ecologically relevant members of the microbiome [44]. Predator-prey co-cultures were prepared as described in Remy et al. in DNB medium (1:10 diluted nutrient broth, supplemented with 2 mM CaCl_2_ and 3 mM MgCl_2_) at pH 7.4 [75]. Liquid co-cultures were incubated at 28°C for 48 h or until the culture cleared, indicating active predation. Clear co-cultures were filtered through a 0.45 µm filter to remove remaining prey cells. These predator filtrates were then spotted onto double-layer DNB agar plates, with a lower layer containing 1.5% agarose and an upper layer containing 0.7% agarose and *E. coli* ML35, as described in Jurkevitch et al. [51]. Both agar phases were prepared at pH 7.4, 6.8, or 6.5. One drop of filtrate was added to the surface of each plate, and plates were incubated inverted at either 28°C or 37°C for up to 10 days. Clearing of *E. coli* within the top agar layer was taken as evidence of predatory activity.

#### Predation in mucin medium

Predation by *B. bacteriovorus* HD100 on prey *E. coli* ML35 in porcine mucin was tested in liquid culture. DNB medium (supplemented with 2 mM CaCl₂ and 3 mM MgCl₂) was prepared with either 2 mg/ml or 5 mg/ml porcine mucin (Sigma, cat. no. M2378). Co-cultures were prepared according to Remy et al. [75], alongside a predator-free control, and incubated at 28°C with shaking at 180 rpm for 48 h. Clearing of the co-culture relative to the control was taken as evidence of predation.

#### Phylogenetic tree of reductases

SwissProt proteins of bacterial nitrite reductases (search terms EC: 1.7.2.1 and taxonomy: bacteria) with the highest evidence score of five were downloaded from Uniprot (N=16), and ten additional lower evidence sequences of potential Bdellovibrio reductases (search terms EC: 1.7.2.1 and taxonomy: *Bdellovibrio*) with unreviewed status. We added four additional nitrite reductase candidates from our own collection of *Bdellovibrio* isolates, whose genomes we previously published [44] and for which nitrite reductases could be annotated using Bakta [76] (v1.9 with database v5 full). The multiple sequence alignment was done with MAFFT [77] (v7.526; settings: –auto), and poorly aligned regions were trimmed with trimAl [78] (v1.5.rev0; settings: -automated1). A tree was then inferred using maximum-likelihood phylogeny estimation with IQ-TREE [79] (v2.3.6; settings: -m MFP -bb 1000 -alrt 1000 -nt AUTO). The single best tree identified by maximum-likelihood optimization was then plotted using ggtree [80] in R.

## Results

To investigate the relevance of BALOs as drivers of microbiome diversity, we conducted controlled experiments using the defined CeMbio 12- and 43-member bacterial communities, which were exposed to *Bdellovibrio* strains with either broad (MYbb2) or narrow (MYbb4) prey ranges. We then identified the prevalence of predatory bacteria in the human microbiome by searching for BALO ASVs in two publicly available datasets. The Human Microbiome Catalog (HMC) [49] provided a large collection of gut-specific samples, while the Phenotypic Reference for Integrated Microbiome Enrichment (PRIME) [50] dataset encompassed all human body sites. Together, these datasets enabled a comprehensive meta-analysis of predatory bacteria across human-associated microbial communities. Next, we investigated the likelihood that BALOs could survive under gut-like conditions. We tested different *Bdellovibrio* strains under conditions relevant to the human gut, including low pH, body temperature, and growth in mucus-supplemented medium. Additionally, we screened available BALO genomes for nitrite reductase genes. Finally, we propose a conceptual framework for how BALOs could contribute to gut health by specifically targeting facultative anaerobes that otherwise could exploit dysbiotic conditions.

### BALO diversity effect & and relevance in human microbiomes

#### Diversity effect in the CeMbio community

We first assessed whether the presence of BALOs affects community diversity, including richness, in line with ecological theory. We performed laboratory experiments in flasks using either a 12-strain community (CeMbio12) or a 43-strain community (CeMbio43), each cultivated with or without a BALO. In addition, we included two BALOs that differ in their prey range: MYbb2, which has a broad prey range (4 out of the 12 CeMbio strains), and MYbb4, which is restricted to bacteria of the genus *Ochrobactrum* (i.e., 1 out of the 12 CeMbio strains) [40].

After 12 days of co-cultivation (with serial transfers every 12 hours), we observed an overall decline in community richness over time (Fig. S2). However, communities exposed to BALOs showed a significantly slower decline in richness than control treatments (Fig. 1A, B, Fig. S2). Community richness ultimately stabilized at a similar level in both BALO-treated and control communities. Notably, however, BALOs were not detected at the end of the experiment, suggesting that their die-out may account for the convergence in richness between treatments.

**Fig. 1:**
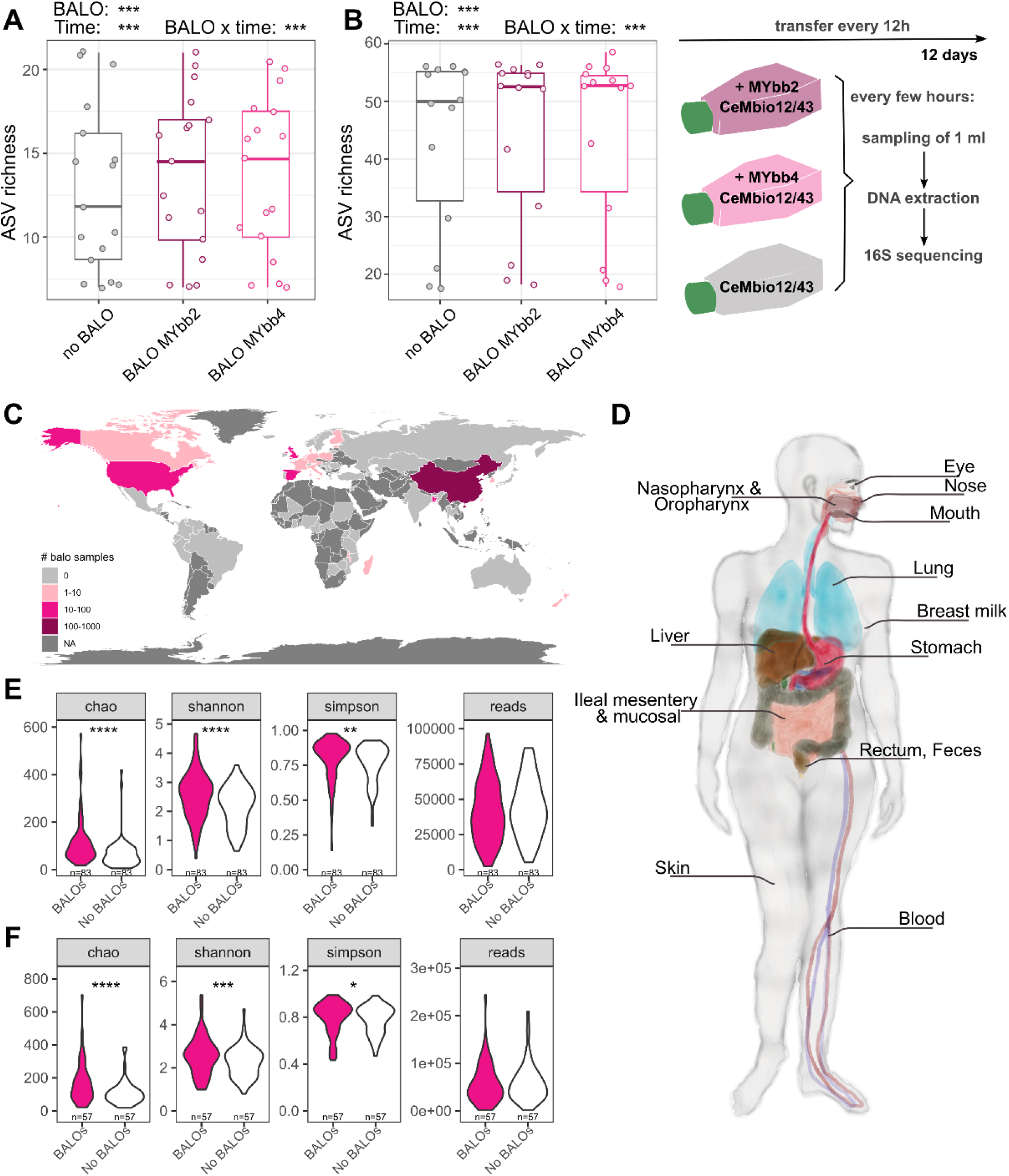
BALO diversity effect and relevance in human microbiomes. **A, B)** BALO presence significantly affects the community richness of CeMbio12 **(A)** and CeMbio43 **(B)** in vitro. Time-resolved data are available in Fig. S2. The ASVs identified as BALOs were excluded from the analysis. Not all shown ASVs mapped to known CeMbio12/43 sequences at ≥99.5% identity, but were kept to account for sequencing errors and/or adaptation. Statistical analysis was performed using a generalized linear mixed model with a Conway-Maxwell-Poisson (CeMbio12) or Gaussian distribution (CeMbio43) over multiple sampling time points (*Richness ∼ BALO × time + (1 | replicate)*). **C)** Global map of BALO-positive samples showing the number of BALO samples found. NA indicates that no study was available covering the country. **D)** Tissue, fluid, or body regions where BALOs were found. **E, F)** Comparison of alpha diversity measures (Chao, Shannon, Simpson) between studies that contained BALOs in **(E)** the HMC and **(F)** the PRIME dataset. Median values of samples with and without BALOs. A paired Wilcoxon test was used. P-values: p ≤ 0.0001: ’****’, 0.0001 < p ≤ 0.001: ’***’, 0.001 < p ≤ 0.01: ’**’, 0.01 < p ≤ 0.05: ’*’.

#### BALOs are prevalent in human microbiomes across the body and worldwide

Using the newly available HMC and PRIME databases, we investigated whether BALOs are present in human microbiomes. Both databases aggregate studies that contain 16S rRNA gene sequencing data from diverse human samples, with sample sizes varying across studies. The shares of samples containing BALOs were 1% for HMC and 3.4% for PRIME. We identified studies containing at least one BALO-positive sample (17% of studies in HMC and 44% in PRIME). In those projects, BALOs were observed at 2.4% (HMC) and 3.9% (PRIME). The PRIME dataset contains samples from all human microbiomes; when restricted to the gastrointestinal system, BALO samples, as a percentage of BALO projects, account for 2.2%, which is very close to the 2.4% found in HMC (Tab. 1).

**Tab. 1:** Details about datasets that were used in our study.

| Dataset | Projects | Samples | BALO projects* | Used samples* | BALO samples | BALO frequency# |
| --- | --- | --- | --- | --- | --- | --- |
| HMC<sup>a</sup> | 492 | 168464 | 83 (17%) | 37377 | 909 | 1% / 2.4% / 2.4% |
| PRIME<sup>b</sup> | 129 | 53449 | 57 (44%) | 46351 | 1809 | 3.4% / 4% / 2.2% |

<sup>a</sup> Human Microbiome Compendium (**HMC**): Abdill2025 Integration of 168,000 samples reveals global patterns of the human gut microbiome. Cell https://doi.org/10.1016/j.cell.2024.12.017
<sup>b</sup> Phenotypic Reference for Integrated Microbiome Enrichment: Zhang2026 **PRIME**: a database for 16S rRNA microbiome data with phenotypic reference and comprehensive metadata. Nucleic Acids Research https://doi.org/10.1093/nar/gkaf1057
\* **BALO project** is a project that has at least one sample with a detected BALO
+ Samples from BALO projects that were used for analysis

BALO-positive samples were not distributed evenly across geographic regions. Leveraging metadata in the HMC, we found that BALO-positive samples originated primarily from North America, China, India, and several European countries (Fig. 1C). This pattern was independent of the overall geographic distribution of samples, suggesting that a high sample volume from a specific country did not necessarily correlate with the presence of BALOs (Fig. S3). For instance, while samples from Germany, Russia, and Australia were numerous, none contained BALOs. Generally, we observed a positive correlation between the total number of samples and the number of BALO-positive samples per region. However, some countries diverged from this trend, appearing either more “BALO-rich” than expected (e.g., China and South Korea) or notably less so (e.g., the Netherlands and Denmark; Fig. S4).

To determine the specific localization of BALOs within the human host, we leveraged detailed metadata from the PRIME database, which includes samples from all human microbiomes and specifies the organ or body site from which each sample was obtained. BALO-positive samples were identified across a wide range of sites throughout the human body (Fig. 1D, Fig. S5). We also observed sites where BALOs were absent; however, not all sites were sampled to the same extent. Interestingly, BALOs were identified in body sites that would seemingly not favor their growth, such as the gut, which is characterized by low pH and oxygen-limited or anaerobic conditions. In particular, BALOs were rarely detected in fecal samples (0.3% of the total) but were more frequent in mucosal biopsies (59% of the total) (Fig. S5). This distribution suggests that BALOs inhabit a specific niche within the gut, likely proximal to the epithelium, where oxygen levels are slightly elevated due to the metabolic activity of host cells.

#### Diversity effect in human microbiomes

To investigate whether the presence of BALOs correlates with alpha diversity in human microbiomes, we focused on studies that included BALO-positive samples. Within these studies, we compared BALO-positive and BALO-negative samples using Chao1, Shannon, and Simpson diversity indices (Fig. 1 E-F).

To account for potential sampling bias, specifically, the increased likelihood of detecting BALOs in samples with higher sequencing depth, we also compared read counts between these groups. Across both the HMC and PRIME datasets, BALO presence was consistently and significantly associated with higher microbiome diversity, while read counts did not differ between BALO-positive and BALO-negative samples. The effect was most visible with Chao1 and less so with Simpson diversity, indicating a richness rather than an evenness effect (Fig. 1 E-F). The findings were consistent across most body sites and geographic regions, with only a few exceptions, such as lung lavage fluid, ileal mesentery, and nasal skin (Fig. S5), as well as sub-Saharan Africa and Central/Southern Asia (Fig. S6).

We next investigated whether the presence of BALOs is associated with changes in overall community structure in human microbiomes. Specifically, we hypothesized that BALO predation reshapes microbial communities by reducing the abundance of dominant taxa, thereby freeing ecological niches for otherwise less competitive or extinction-prone species. Under this framework, communities containing BALOs would be expected to be less centralized and rely less on highly connected key nodes. To test this, we constructed co-occurrence networks from BALO-free and BALO-containing samples. When considering all significant edges, the BALO-free dataset yielded a larger, more highly connected network, comprising nearly 250,000 edges compared to ∼185,000 in the BALO-containing network (Fig. S7). However, this difference is likely influenced by the substantially larger sample size of the BALO-free dataset (55,287 samples vs. 1,110 samples). After filtering out weak interactions, the network derived from BALO-containing samples became comparatively larger, suggesting that many edges in the BALO-free network represent weak, albeit statistically significant, associations (Fig. S8). Overall, the BALO-containing network was less dense and less centralized and exhibited a higher proportion of negative edges (Fig. S7, Tab. S1), consistent with a potential role of predation in structuring microbial interactions. Interestingly, the BALO-associated network included one node belonging to the Oligoflexia, a group within the Bdellovibrionota. However, as this lineage has not yet been cultivated, its predatory capacity remains uncertain, and it was not included in our general diversity analysis.

#### BALOs in gut microbiomes

Next, we investigated whether the positive correlation between BALO presence and microbial diversity holds true for both healthy and diseased individuals. We focused specifically on human gut microbiomes, as gut-related pathologies are frequently characterized by dysbiosis, typically involving an enrichment of facultative anaerobes and reduced alpha diversity. Since the majority of tested BALO prey species belong to facultative anaerobes (74%, Tab. S2), we hypothesized that BALOs might play a regulatory role during disease, when their potential prey bacteria are most abundant. Our results showed that the presence of BALOs remains significantly associated with samples from different parts of the gastrointestinal (GI) tract (e.g., stomach, ileum, stool; Fig. 2A; for all body sites, cf. Fig. S5). Although most samples were from stool, those showed a low BALO prevalence (< 1%). In contrast, other sites, such as ileal mucosa, contained more BALO than non-BALO samples, consistent with the reported underestimation of facultative bacteria in feces compared with rectal swabs or biopsies [81]. Upon closer examination of specific clinical conditions, we found that the presence of BALOs in GI tract samples correlates strongly with increased Chao1 diversity in patients with, e.g., IBD, heart failure, and gastric adenocarcinoma (Fig. 2B; for diseases from samples across all body sites, cf. Fig. S9). The association observed in IBD patients is particularly striking, as this condition is typically characterized by reduced microbiome diversity, as confirmed by our data analysis (Fig. 2D), and systemic inflammation. Inflammatory processes often lead to the accumulation of reactive oxygen species (ROS) and increased local oxygen availability, which may favor BALO activity. Specifically, the rapid, flagellum-driven motility of these predators is energetically demanding and likely benefits from the enhanced metabolic potential offered by oxygenated environments. Thus, under IBD conditions, BALOs may encounter an expanded ecological niche characterized by increased oxygen availability and a high abundance of prey, as the facultative anaerobes that proliferate during inflammation could serve as primary prey.

**Figure 2:**
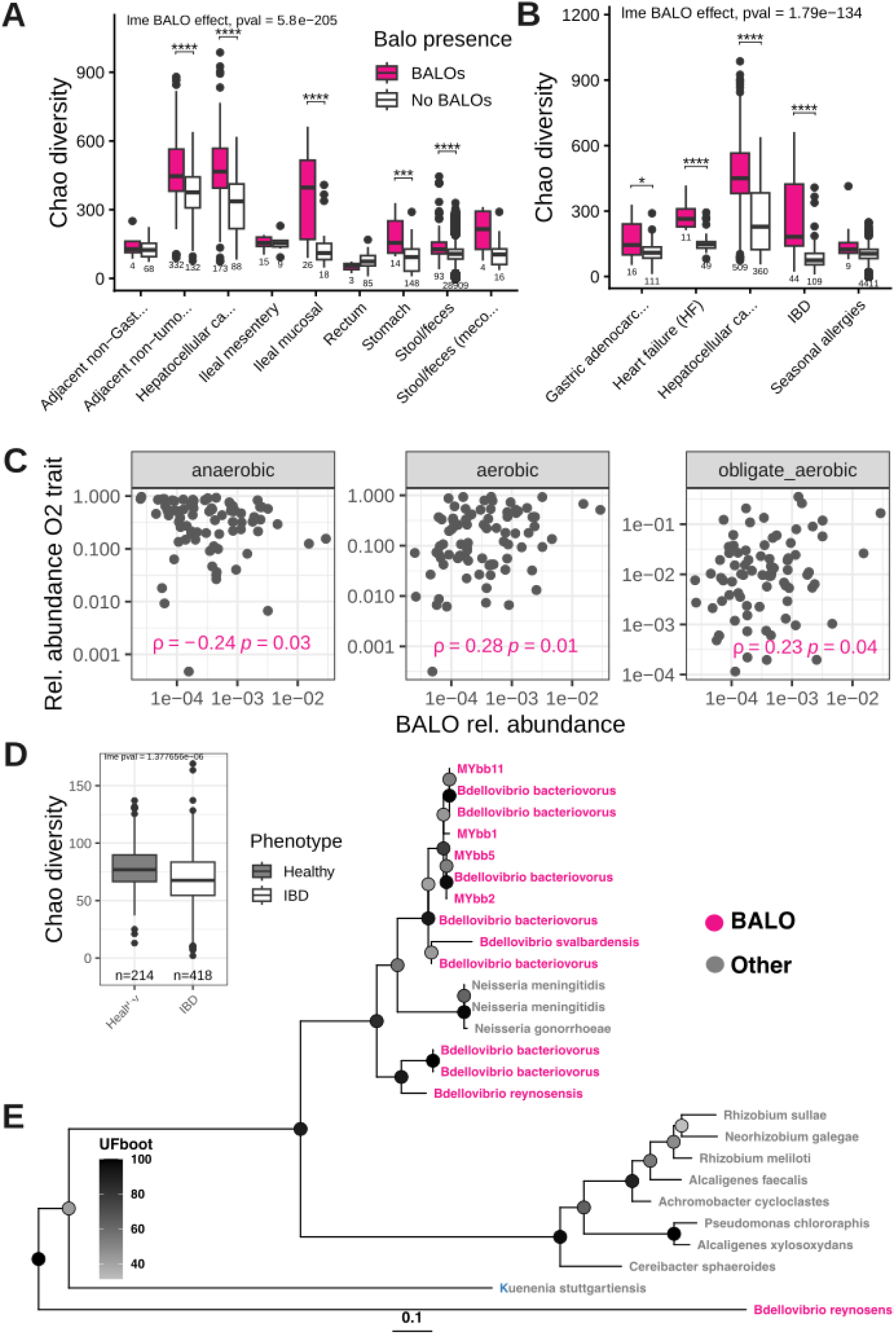
BALOs in human gut microbiomes. Chao alpha diversity of gut samples across gut body sites and diseases **(A, B)**. Samples from the PRIME dataset were selected from the “Digestive System/Gut” system and shown according to **(A)** body site with at least two BALO samples and **(B)** diseases with at least five BALO samples. The exact number of samples for each category is shown below the boxplot. The Wilcoxon test was performed to compare samples with and without BALO; p-values were adjusted using the BH method. A linear mixed-effects model that included study and total read count as random effects confirmed the positive impact of BALOs on alpha diversity. **C)** Spearman correlation of relative abundance of oxygen preferences aggregated for each sample from species traits correlating with BALO abundance. The comparison is between the median abundance of BALO samples for each study (n=83) of the HMC dataset. The oxygen preference per species is predicted from the 16S sequence using AmpliconTraits[70], and the values are summed for each sample. Only oxygen traits with significant correlation were shown (cf. Fig. S13 for all traits). **D)** Alpha diversity of IBD samples. Studies from the PRIME dataset with IBD samples and corresponding healthy controls (4 studies in total). Linear mixed-effect model to account for read depths and study impact. **E)** Phylogenetic tree of nitrite reductases from reference species and BALO species. Reference sequences were obtained from SwissProt (EC number 1.7.2.1), and the tree was created using iqtree. P-values: p ≤ 0.0001: ’****’, 0.0001 < p ≤ 0.001: ’***’, 0.001 < p ≤ 0.01: ’**’, 0.01 < p ≤ 0.05: ’*’. Abbreviations: “Stool/feces (meco…” = Stool/feces (meconium), “Adjacent non-Gast…” = Adjacent non-Gastric adenocarcinoma tissue liver tissue (hepatocellular carcinoma, HCC), “Hepatocellular ca…” = Hepatocellular carcinoma tissue (HCC, liver), “Adjacent non-tumo…” = Adjacent non-tumor liver tissue (hepatocellular carcinoma, HCC), “Hepatocellular ca…” = Hepatocellular carcinoma (HCC), “Gastric adenocarc…” = Gastric adenocarcinoma.

Next, we examined whether BALOs occur more frequently in samples from healthy or diseased individuals. Based on our earlier prediction, we expected a higher prevalence of BALOs in diseased microbiomes, where enrichment of potential prey bacteria would create a more favorable environment for predators and where their predatory activity could provide “probiotic” benefits by mitigating dysbiosis. Supporting this, we observed that the frequency of BALO-positive samples was significantly higher in disease-associated microbiomes compared to healthy controls (Fig. S10). To clarify the potential health effects, we further screened for studies that contained healthy controls with BALO samples. Here, we found eight studies on six diseases, two of which specifically included gut samples. Across most diseases, BALO presence was significantly and positively associated with higher alpha diversity, with sample count as a limiting factor, for example, in IBD (Fig. S11).

#### How can BALOs survive in the GI tract?

Finally, we sought to more clearly define the ecological niche that BALOs likely occupy in the human gut. Our earlier results indicate that BALOs can survive the low-oxygen conditions characteristic of the human intestine. To corroborate this, we leveraged genome-based predictions to assess the oxygen preferences of microbial communities that were either BALO-free or BALO-positive.

Consistent with their known physiology, the presence of BALOs was significantly associated with a (micro)aerobic lifestyle and negatively correlated with an anaerobic lifestyle (Fig. 2C, Fig. S12, Fig. S13). Simultaneously, we observed a positive trend linking BALOs to the presence of facultative anaerobes. Although this specific association did not reach statistical significance, it is important to note that a facultative anaerobic lifestyle is generally more difficult to predict accurately from genomic data than an obligate lifestyle. In addition, many aerobes are potential facultatives. This supports our hypothesis that BALOs share a niche with these bacteria, which in most cases (>70%) serve as their primary prey, as evidenced by our literature screening (Tab. S2).

We next directly tested the physiological growth conditions of several BALO strains, including the commonly used strain *B. bacteriovorus* HD100, which was recently recovered from the human gut [47], or, more precisely, a closely related isolate thereof, indicating its ability to persist under gut-like conditions. Together with two strains isolated from the direct environment of the nematode *Caenorhabditis elegans* (and frequently found in *C. elegans* microbiome data; MYbb2 and MYbb1), we found that these strains were capable of active predation at 37 °C and pH 6.5 (plaque formation). In contrast, two other *C. elegans*-derived strains (MYbb7 and MYbb5) were active at pH 6.5, but not at 37 °C. We then tested whether predation by *B. bacteriovorus* HD100 can occur in a medium containing mucin (porcine gastric mucin) to approximate conditions near the gut epithelium. Indeed, we found that HD100 can infect *E. coli* ML35 in medium supplemented with either 2 mg/ml or 5 mg/ml porcine mucin (Fig. S14). These results suggest that at least a subset of BALOs is capable of active predation under conditions resembling those of the (human) gut.

Finally, we investigated whether BALOs can survive or even persist under anaerobic conditions, despite their association with aerobic lifestyles in our earlier analyses. Such a capability would allow these predators to endure low-oxygen conditions in the gut and potentially become active again when oxygen levels increase, for example, during inflammation. Alternatively, the ability to use electron acceptors other than oxygen would substantially expand their ecological niche. To address this, we analyzed published BALO genomes for the presence of genes involved in anaerobic respiration. Specifically, we screened for nitrite reductases and found that these genes are present in multiple BALO genomes (Fig. 2E). For example, *B. bacteriovorus* HD100 encodes several nitrite reductases and nitric oxide reductases. In previous experiments, it has been shown that *B. bacteriovorus* HD100 can perform predation under anoxic conditions using nitrate, albeit with lower efficiency [82]. Our genome-based analysis, however, indicates that this capacity might be a trait of many *Bdellovibrio* species.

These findings suggest that BALOs may be capable of partial denitrification and/or detoxification of reactive nitrogen species, which are likely encountered in the gut environment, thereby supporting their persistence under oxygen-limited conditions.

Together, our results indicate that BALOs are consistent members of human microbiomes and are also present in the gut, where they predominantly occur at sites with relatively high availability of electron acceptors. This likely enables them to actively prey on Gram-negative bacteria, including facultative anaerobes that are commonly enriched in inflamed gut environments. Through this combined predatory (“probiotic-like”) and antibacterial activity, BALOs may help reshape dysbiotic microbiomes. As such, they represent a promising candidate for future therapeutic strategies aimed at restoring microbial balance in disease-associated states (Fig. 3).

**Figure 3:**
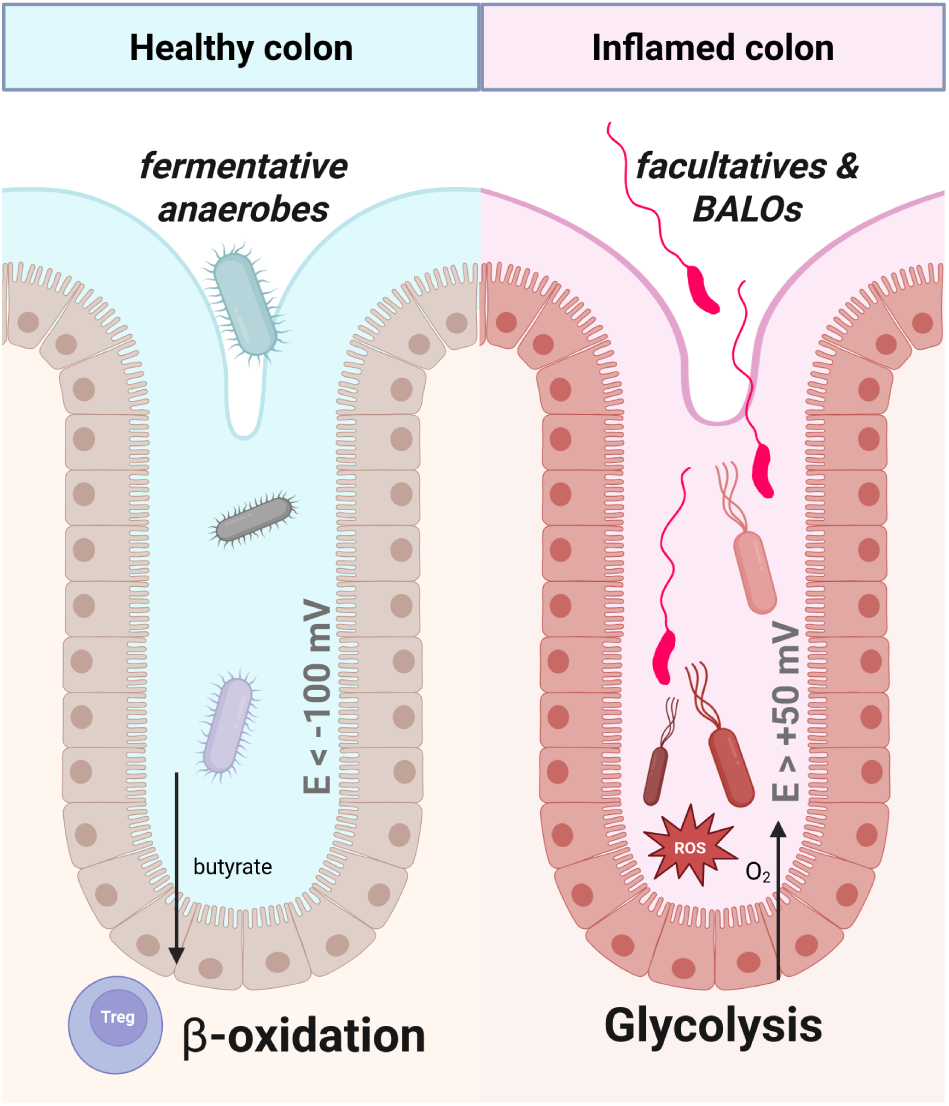
Conceptual framework of how BALOs could contribute to gut health. We found BALOs to be prevalent in human gut microbiomes and associated with dysbiotic conditions in which their prey (facultative anaerobes) could thrive. In addition, they seem to have specific adaptations that enable them to survive in the gut environment (temperature, pH, mucus, and anaerobic respiration). Based on this, we propose that BALOs could enrich under inflammatory conditions (right side of the figure, in red) and specifically target bacteria that exploit these dysbiotic conditions. By inhibiting the dysbiotic overgrowth of facultative anaerobes, BALOs could help maintain fermentative organisms that produce anti-inflammatory compounds such as butyrate. After the inflammation subsides, the niche electron acceptor disappears, and BALOs find less food and decrease in abundance. The redox potential (E) indicates the availability of oxygen (E > 50mV) and fermentative conditions (E < -100 mV). Epithelial cells are known to be polarized between oxidative phosphorylation fueled by beta-oxidation of microbial-produced butyrate (healthy state) and glycolysis with no or reduced respiration and increased oxygen diffusion (diseased state) [83]. In addition, regulatory T-cell (Treg) differentiation is induced by butyrate, further maintaining a non-inflammatory state [84].

## Discussion

Predation shapes biological communities across all kingdoms of life. In bacterial ecosystems, bacterivorous organisms, including bacteriophages, protists, and predatory bacteria, exert top-down control that can free niches occupied by dominant species. The extent of the effect may vary with the predator’s prey range [40, 45]. Here, we focus specifically on predatory bacteria belonging to the BALOs, as they are readily detectable in 16S amplicon data, possess a broad prey range, and have demonstrated potential to shape microbiome composition [38]. Furthermore, BALOs remain largely understudied, particularly regarding their role in human microbiome restoration. Our findings highlight BALOs as significant yet underappreciated drivers of human microbiome dynamics.

Consistent with ecological theory [85–87], our in vitro experiments showed that BALOs maintain microbiome diversity over time by slowing species loss. This is confirmed in our meta-analysis, where the presence of BALOs correlates positively with alpha diversity across human body sites. Network analysis reinforces this pattern: BALO-free networks were centralized and dense, dominated by a few hub taxa and potentially vulnerable to their collapse, whereas BALO-containing networks were decentralized and sparser. By suppressing dominant species, BALO predation appears to distribute stability more broadly across community members.

Given the importance of alpha diversity for human microbiome function [20, 88], identifying strategies to enhance it is a key clinical priority. Probiotics are the most widely marketed and used intervention in this context, yet a recent meta-analysis has questioned their overall benefit in diversity, particularly in healthy microbiomes [89]. BALOs may offer a complementary or alternative therapeutic avenue, acting not by adding biomass but by rebalancing community structure through targeted predation. A long-standing assumption, stemming from their classification as obligate aerobes [90, 91], has been that BALOs are irrelevant to the human gut. Previous studies reported microaerobic survival and growth of BALOs [36, 92] and found them only occasionally in human microbiomes [41, 93], absent from fecal samples but more prevalent in mucosal biopsies of healthy individuals [41]. In our meta-analysis, however, we detected BALOs in fecal samples and found them in the microbiomes not only of healthy humans but even more frequently of diseased humans. This pattern matches the differential abundance of their facultative anaerobic prey: they are more prevalent in microbiomes associated with gut-related diseases [46]. In agreement, we detect a substantially higher prevalence of BALOs in mucosal samples, suggesting a niche proximal to the epithelium. Additionally, we demonstrated that *B. bacteriovorus* HD100 can survive and actively prey in porcine mucus. This is consistent with earlier observations that BALOs can prey in polyvinylpyrrolidone media of increasing viscosity [92] and that HD100 exhibits improved attack rates in a polyethylene glycol medium at 5.4 mPa·s, alongside increased predator swimming speeds [94]. However, the latter study showed that further increasing viscosity negatively affected predation efficiency [94]. The direct relevance of these viscosity ranges to mucus nonetheless remains unclear, given that mucus is a viscoelastic gel whose effective microscale viscosity differs substantially from bulk rheological measurements. Moreover, predation performance in viscous environments may be strain-specific for BALOs, as flagellar architectures likely differ in their suitability for swimming under such conditions [44]. Alternatively, BALOs in gel-like environments may preferentially rely on gliding motility to locate prey, a strategy that may be better suited to surface-associated conditions, as demonstrated in biofilms [95]. Whether either mechanism operates in mucus remains to be directly tested.

This proposed niche is particularly interesting in light of evidence that bacteriophages also inhabit the mucus layer across hosts, adhering to mucin glycoproteins and thereby increasing their probability of encountering and infecting bacteria traversing the mucus matrix [96].

Interestingly, not only BALOs but also facultative anaerobes are better captured by biopsies and rectal swabs [81]. A plausible explanation is that although the gut lumen is largely anoxic, localized microenvironments near the mucosa, or those arising during inflammation, experience transient spikes in electron acceptor availability via epithelial oxygen leakage and immune-derived reactive oxygen and nitrogen species [48]. BALO metabolism appears well-suited to these conditions. Their flagellum-driven motility usually implies an energy-rich aerobic lifestyle, while their genomic capacity for nitrogen respiration (e.g., nitrite and nitric oxide reductases) suggests persistence and potential predation under anoxic settings. In line with this, a previous study showed that HD100 can actively predate under anoxic conditions, but only if nitrate is added [82]. As a terminal electron acceptor, nitrate allows almost as high an energy gain as O_2_, and nitrate respiration is widespread among facultative anaerobes [97]. The emergence of nitrate in the gut has been described in diseased conditions, in which intestinal inflammation produces nitrate from host-produced nitric oxide [98]. Most importantly, the same inflammatory conditions that generate these nitrate niches also favor facultative anaerobes such as *Escherichia coli*, *Salmonella,* and *Klebsiella* [46, 99, 100], all of which are classic BALO prey (Tab. S2). The inflamed gut may therefore simultaneously create habitat and prey for BALOs, as summarized in our conceptual framework (Fig. 3). Here, we propose a potential boom-and-bust dynamic in which BALOs track the expansion of facultative anaerobes during dysbiosis. Facultative blooms are implicated in IBD, diabetes, colorectal cancer, and chronic kidney disease [46]. Enrichment of facultative anaerobes and toxic aerobic conditions lead to a further decline in butyrate-producing fermenting organisms, which reinforces inflammation through colonocyte and immune feedback loops [83, 101]. Because BALOs specifically target facultative anaerobes and self-limit as prey decline, they represent a promising candidate intervention for restoring homeostasis in inflammatory disease.

Finally, our experiments show that specific BALO strains tolerate gut-like conditions, including elevated temperature and low pH. Taken together with the recent isolation of a BALO from human stool [47], these findings argue that BALOs are functional gut inhabitants rather than incidental transients and are promising natural candidates for therapeutic strategies targeting chronic gut disease.

## Conclusion

Together, our findings establish BALOs as consistent and functionally relevant members of the human microbiome. Their positive association with microbial diversity across body sites and disease contexts, combined with their capacity to persist under gut-relevant conditions, points to the mucosal interface as a central arena for predator-prey interactions. By preferentially targeting facultative anaerobes, BALOs are well positioned to counteract dysbiotic blooms and support the restoration of fermentative homeostasis, thereby extending their therapeutic potential as “living antibiotics” far beyond acute infection and antimicrobial resistance into the broader landscape of chronic inflammatory and metabolic disease.

Given growing evidence that microbial predation occurs along a continuum rather than as a rare exception [102], the ecological and clinical impact of microbial predation is likely even greater than current detection methods suggest, warranting its integration into future frameworks for microbiome function and therapy.

## Data Availability

Raw sequencing data generated in this study will be made available in the NCBI Sequence Read Archive (SRA) under the BioProject accession number **PRJNA1481554**. The original files from the HMC and PRIME databases, as well as the extracted and filtered data used for our analysis, can be accessed on **Zenodo** (https://doi.org/10.5281/zenodo.21135573). Lastly, the source code to reproduce the analysis and figures is available on **GitHub** (https://github.com/jotech/balo-microbiome).

## Acknowledgements

We would like to thank Katharina Ratjen and Paul Schreiber for their help with the experiments. Further, we thank the group of Evolutionary Ecology and Genetics (Kiel) for valuable discussions. We thank the Competence Centre for Genomic Analysis (CCGA) Kiel for 16S sequencing of our samples from the in vitro diversity experiment. We thank the authors of the HMC and PRIME databases for providing their resources and support.

## Author contributions

JZ performed the meta-analysis (data analysis and statistics), including the wordmap and oxygen preference predictions. JZ also analyzed published BALO genomes for reductases and generated the tree. JJ performed the laboratory experiments and performed the corresponding data analysis and statistical analysis and did the network analysis from HMC data. Both authors conceptualized the study, wrote the draft manuscript and revised and approved the final version of the text.

