## Supplementary figures and tables for "Predatory bacteria as members of human microbiomes and their impact on gut diversity and homeostasis"

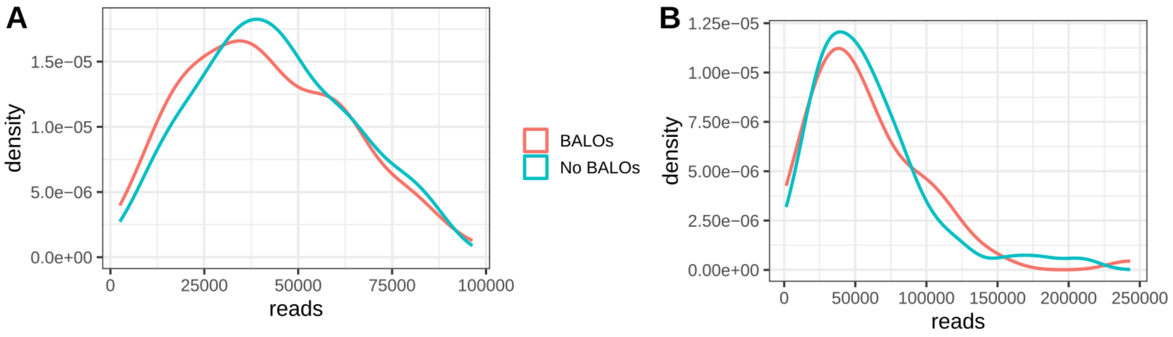


**Fig. S1: Density plot of read counts of BALO and no-BALO samples for HMC (A) and PRIME (B).** Distribution is based on the median values of BALO and no-BALO samples per project.

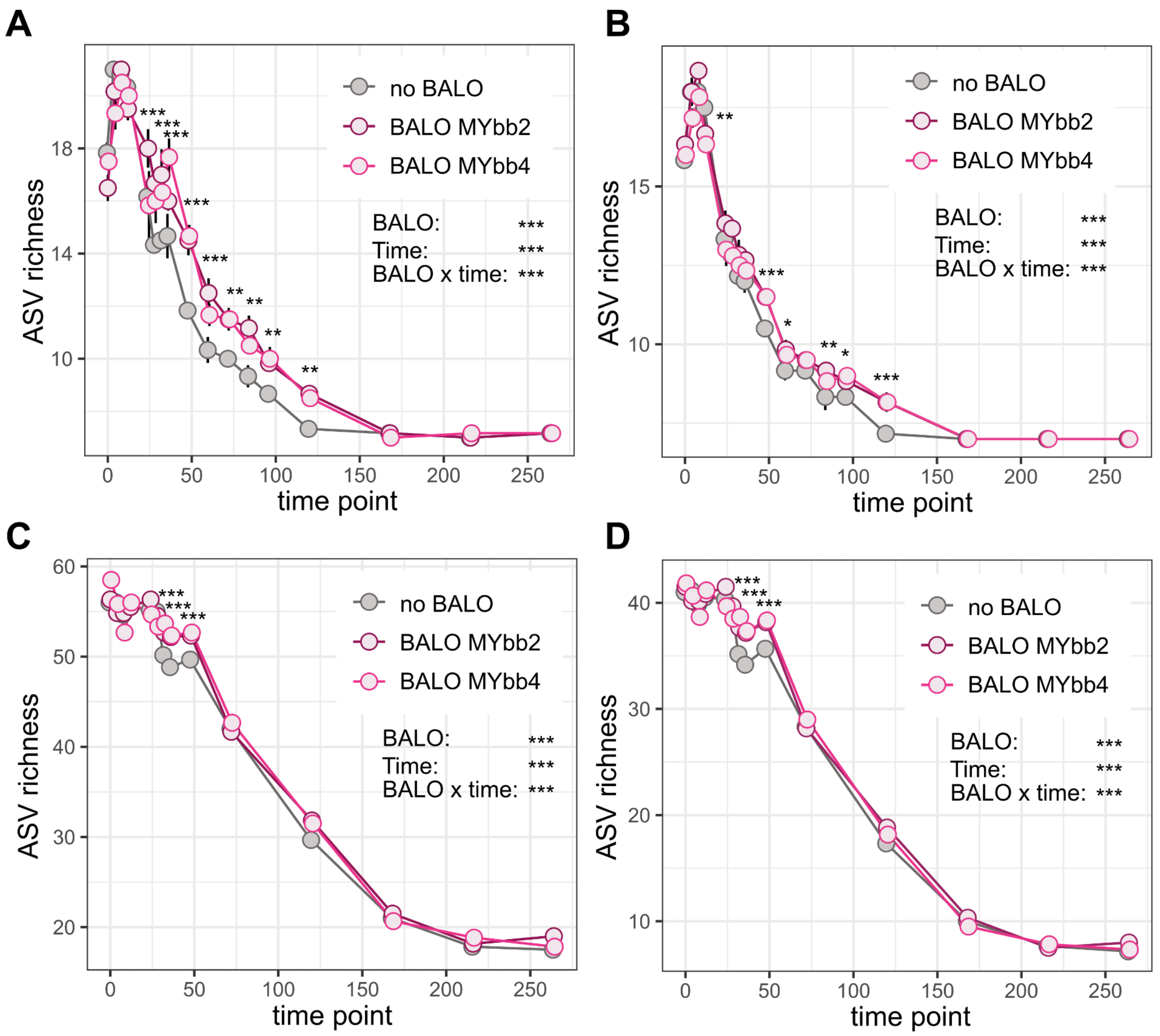


**Fig. S2: BALO presence significantly slows the decline in CeMbio community richness *in vitro*.** The ASVs identified as BALOs were excluded from the analysis. Not all shown ASVs mapped to known CeMbio12/43 sequences at ≥99.5% identity but were either kept to account for sequencing errors and/or adaptation **(A and C)** or removed **(B and D)**. Error bars represent the standard error of the mean across biological replicates. Colors indicate treatment groups: no BALO (gray), broad prey-range BALO MYbb2 (purple), and narrow prey-range BALO MYbb4 (pink). Statistical analysis was performed using a generalized linear mixed model with a Conway-Maxwell-Poisson (COM-Poisson) distribution to account for underdispersion (*Observed ~ BALO × time + (1 | replicate)*) in A and B and a gaussian distribution in C-D, followed by pairwise comparisons using estimated marginal means (emmeans) with Benjamini–Hochberg correction for multiple testing. P-values: $p \leq 0.0001$: '****', $0.0001 < p \leq 0.001$: '***', $0.001 < p \leq 0.01$: '**', $0.01 < p \leq 0.05$: '*'.

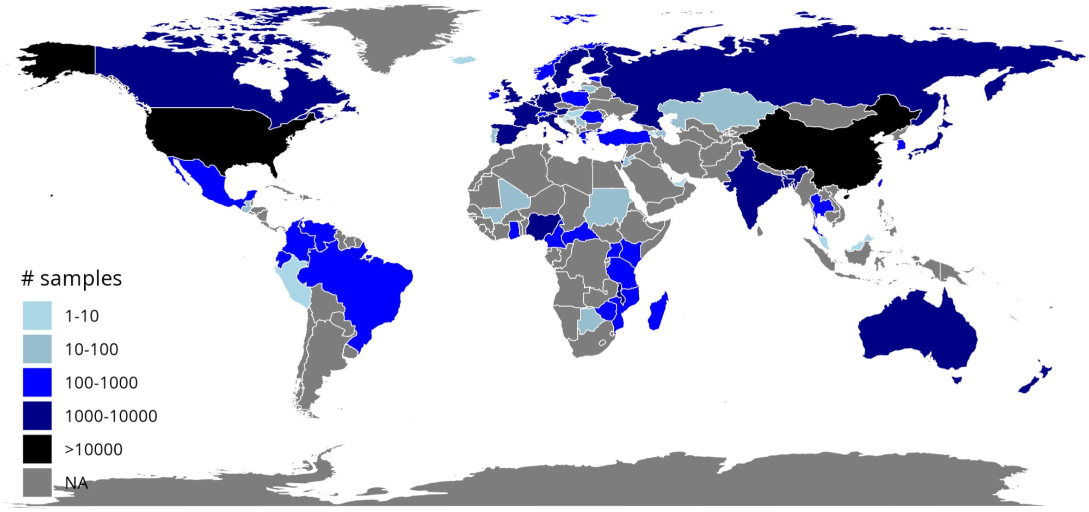


**Fig. S3: World map with the number of samples per country.** NA indicates that no study was available covering the country.

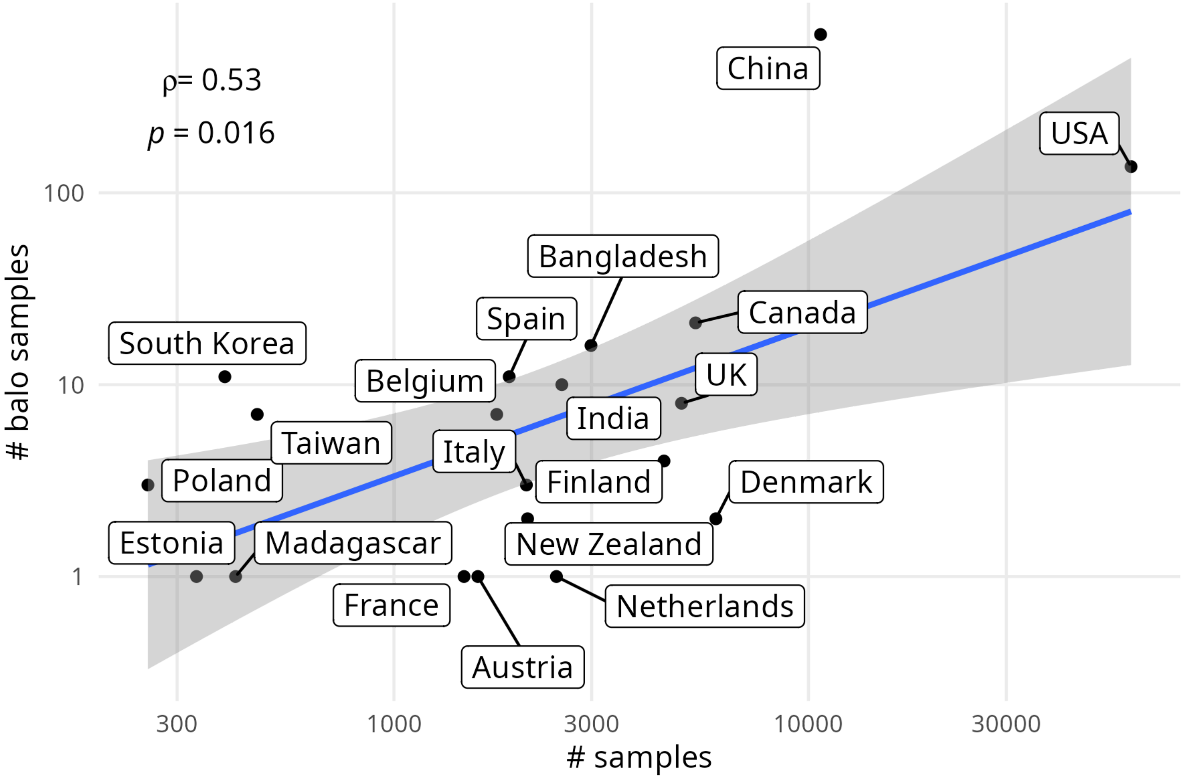


**Fig. S4: Total number of samples vs. samples with BALO reads per country.** Spearman correlation and linear regression with confidence intervals are shown.

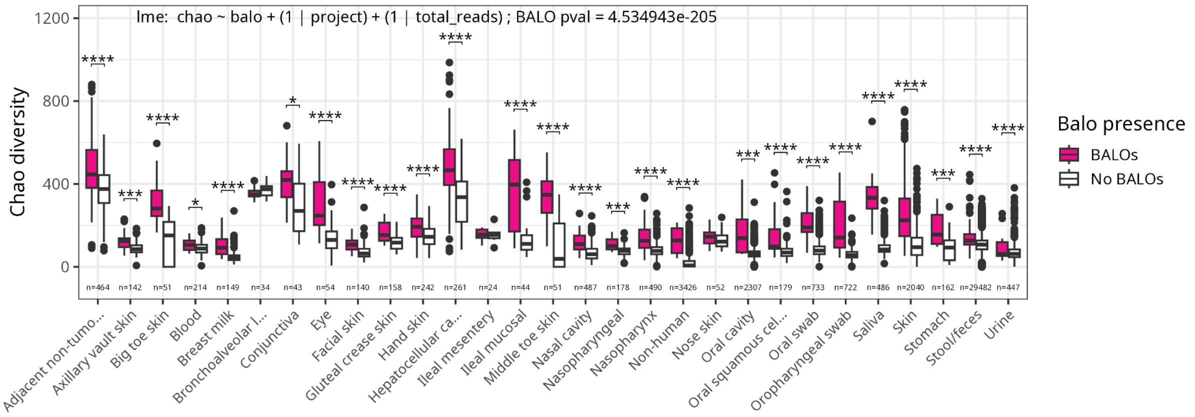


**Fig. S5**: Chao alpha diversity of samples with and without BALOs across body sites in the PRIME dataset. Only body sites with at least 10 BALO samples are shown. The Wilcoxon test was performed to compare samples with and without BALO. A linear mixed-effects model that included study and total read count as random effects confirmed the positive impact of BALOs on alpha diversity. P-values: p ≤ 0.0001: '****', 0.0001 < p ≤ 0.001: '***', 0.001 < p ≤ 0.01: '**', 0.01 < p ≤ 0.05: '*'.

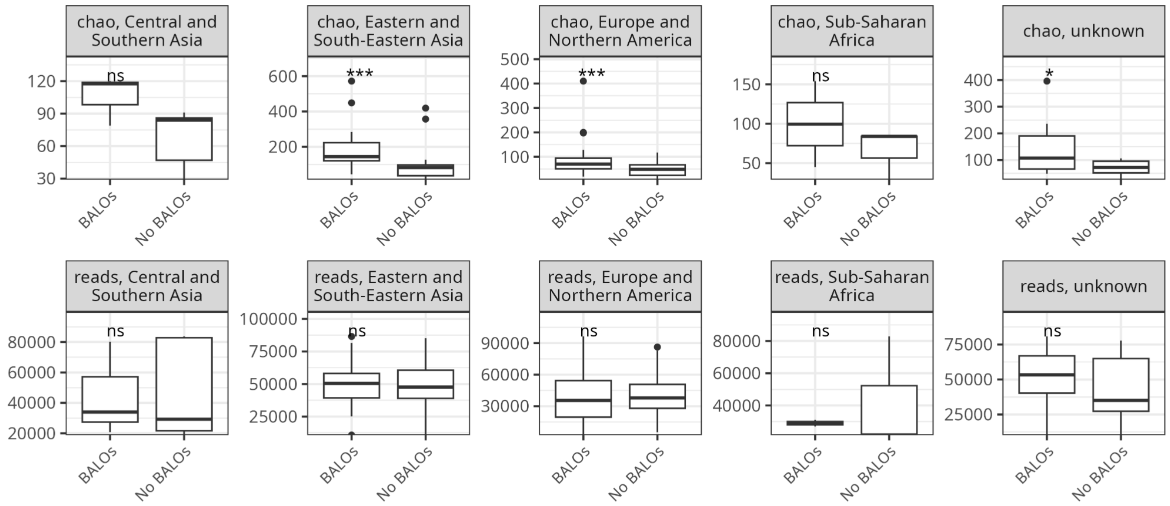


**Fig. S6: Mean alpha diversity and reads of BALO projects per region.** P-values: p ≤ 0.0001: '****', 0.0001 < p ≤ 0.001: '***', 0.001 < p ≤ 0.01: '**', 0.01 < p ≤ 0.05: '*'.

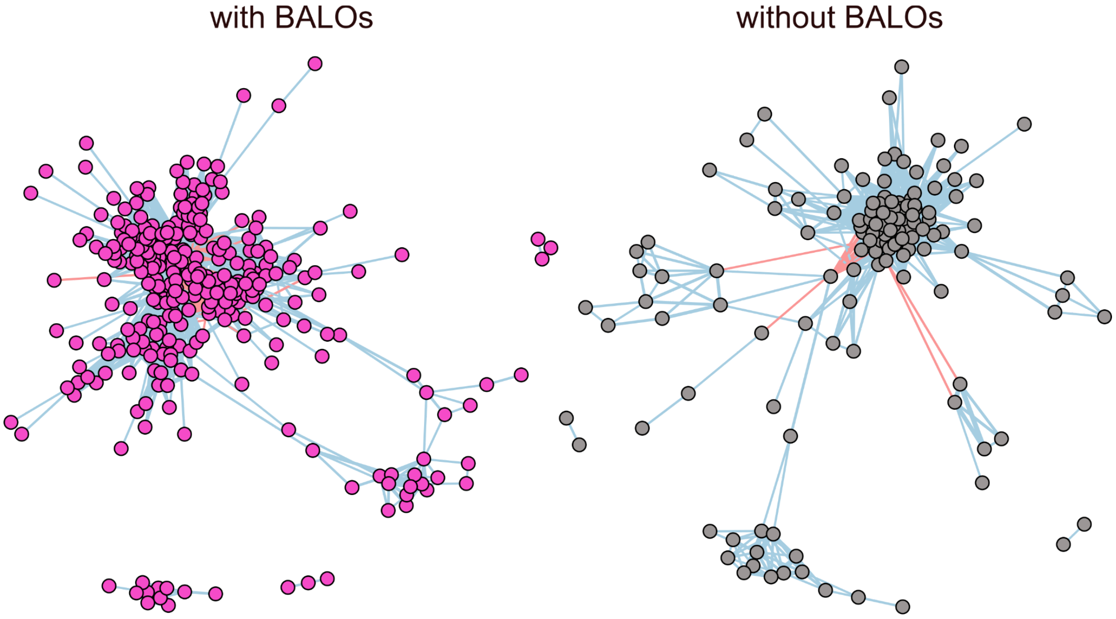


**Fig. S7: Networks generated from samples with BALO presence and samples without BALO presence.** Circles represent nodes (i.e. ASVs). Lines between circles represent edges (i.e. a significant correlation) between two nodes. Only significant edges ($p > 0.05$) and a mean correlation of 0.3 were considered. Blue edges indicate positive correlations while red edges represent negative correlations.

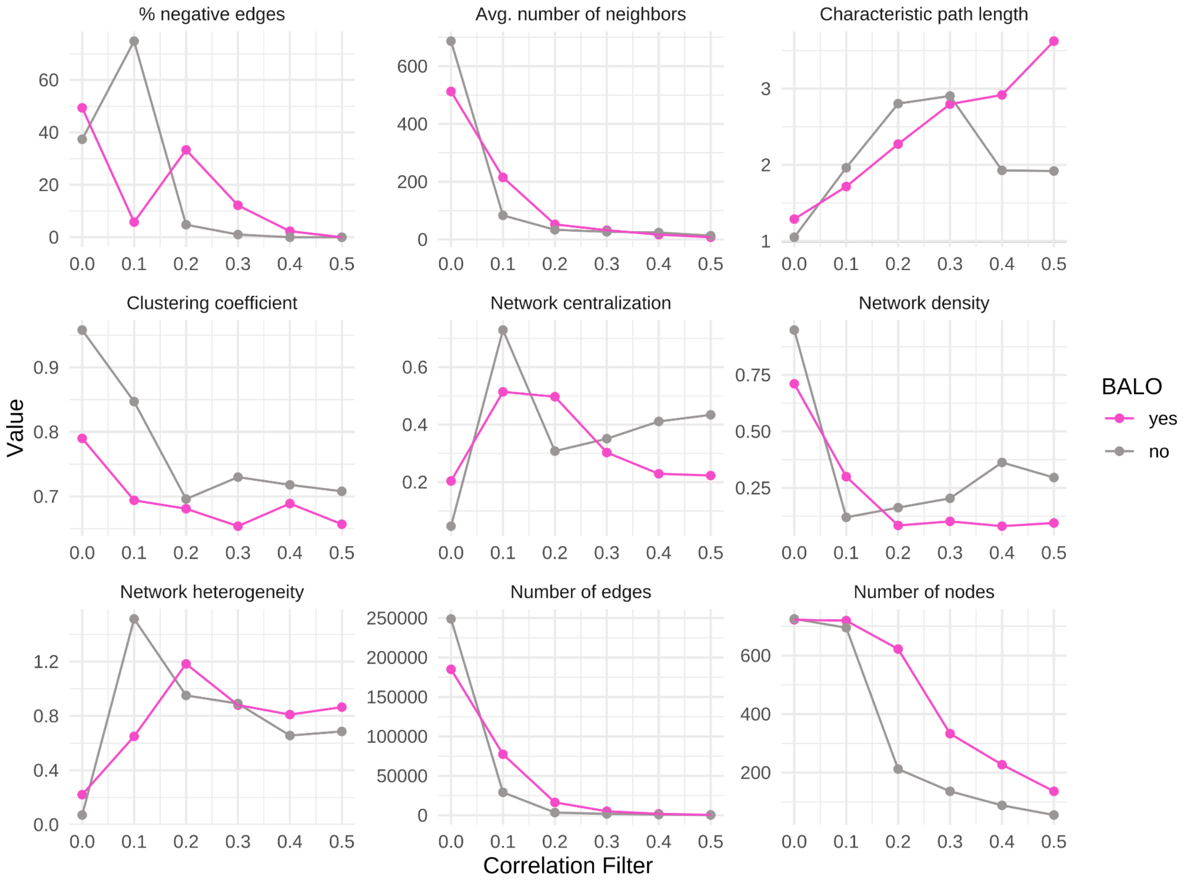


**Fig. S8: Different characteristics of networks change with increasing correlation filtering**. An increasing correlation threshold (up to 0.5) was set to filter edges between significantly correlated ASVs from the network containing BALO-true samples (purple) and BALO-false samples (green).

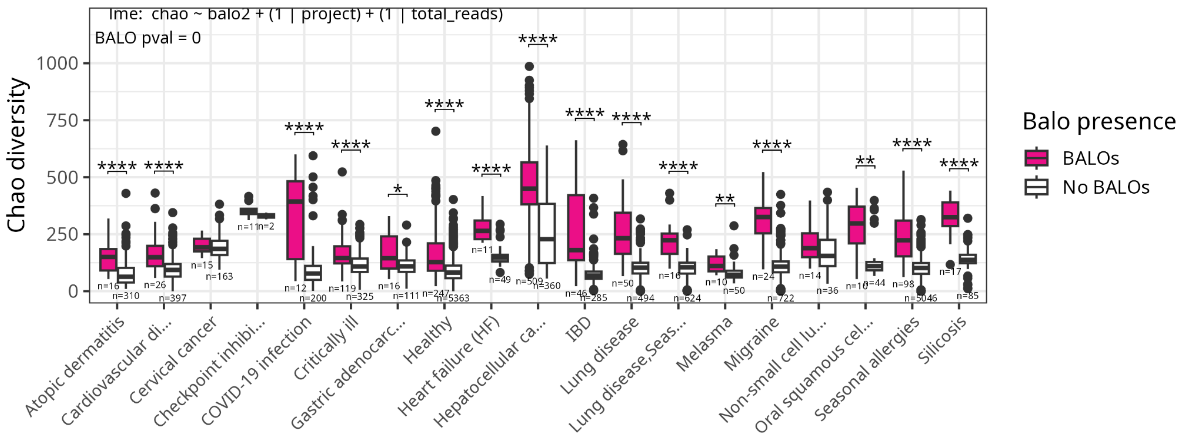


**Fig. S9**: Chao alpha diversity of samples with and without BALOs across diseases in the PRIME dataset. Only diseases with at least 10 BALO samples are shown. The Wilcoxon test was performed to compare samples with and without BALO. A linear mixed-effects model that included study and total read count as random effects confirmed the positive impact of BALOs on alpha diversity. P-values: p ≤ 0.0001: '****', 0.0001 < p ≤ 0.001: '***', 0.001 < p ≤ 0.01: '**', 0.01 < p ≤ 0.05: '*'.

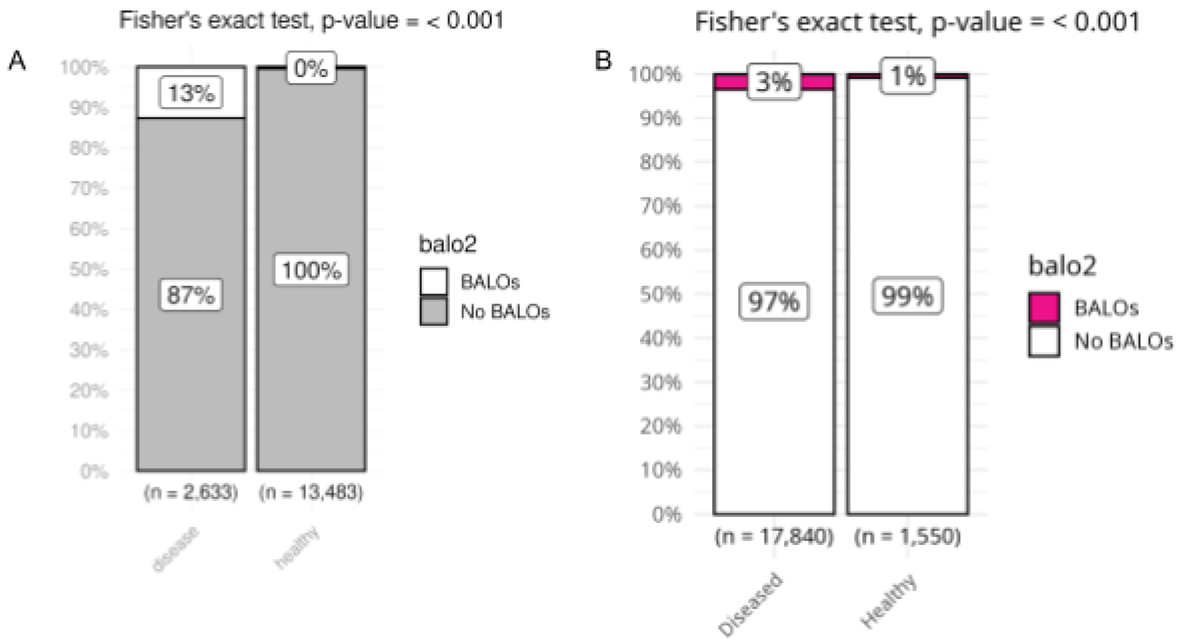


**Fig. S10: Comparison of the frequency of samples containing BALOs in healthy and diseased samples for the A) HMC and B) PRIME databases.** Fisher's exact test was used to assess the likelihood based on the contingency table.

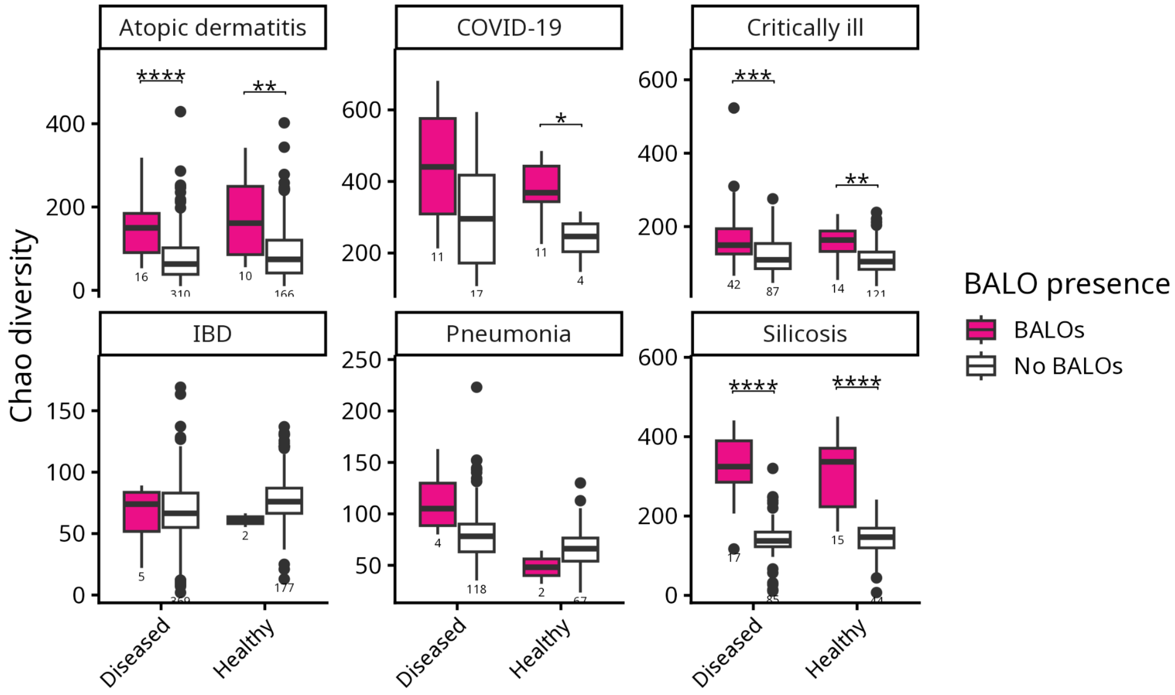


**Fig. S11: Comparison of alpha diversity (Chao) between healthy and diseased samples that contained BALOs.** Studies of the PRIME dataset were included if they contained BALOs in diseased and healthy samples, resulting in eight studies. Studies were grouped according to diseases. The Wilcoxon test was used for significance testing, and p-values were adjusted using the BH method. A linear mixed-effect model with BALOs, disease type, and health status as fixed effects, and total reads and study as random effects, confirmed a significant effect of BALO presence <2e-16. Samples from “Atopic dermatitis” and “Critically ill” were taken from skin, “COVID-19” samples from conjunctiva (eye), IBD and silicosis from GI tract, and pneumonia samples from nasopharynx (nose). P-values: $p \leq 0.0001$: '****', $0.0001 < p \leq 0.001$: '***', $0.001 < p \leq 0.01$: '**', $0.01 < p \leq 0.05$: '*'.

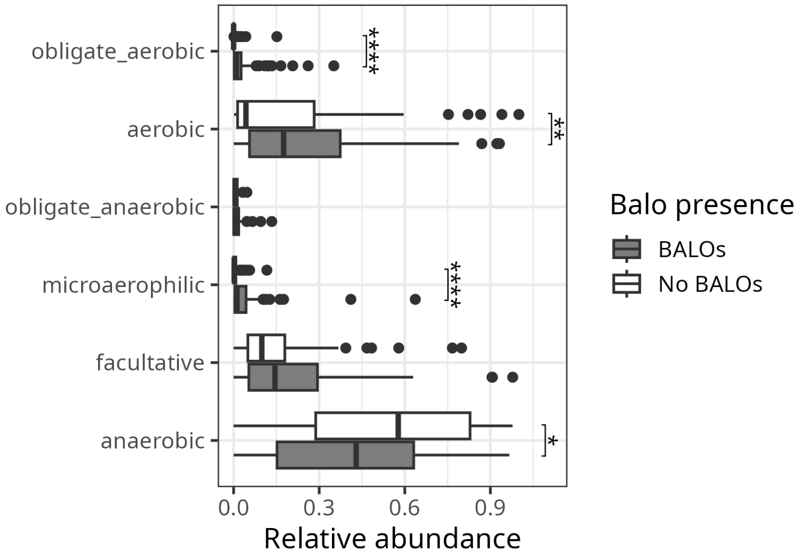


**Fig. S12: Relative abundance of oxygen preferences aggregated for each sample from species traits.** The comparison is between the medians of BALO and non-BALO samples for each study (n=83) of the HMC dataset. The oxygen preference per species is predicted from the 16S sequence using AmpliconTraits. P-values: p<0.0001:'****', 0.0001>p<0.001:'***', 0.001>p<0.01:'**', 0.01>p<0.05:'*'.

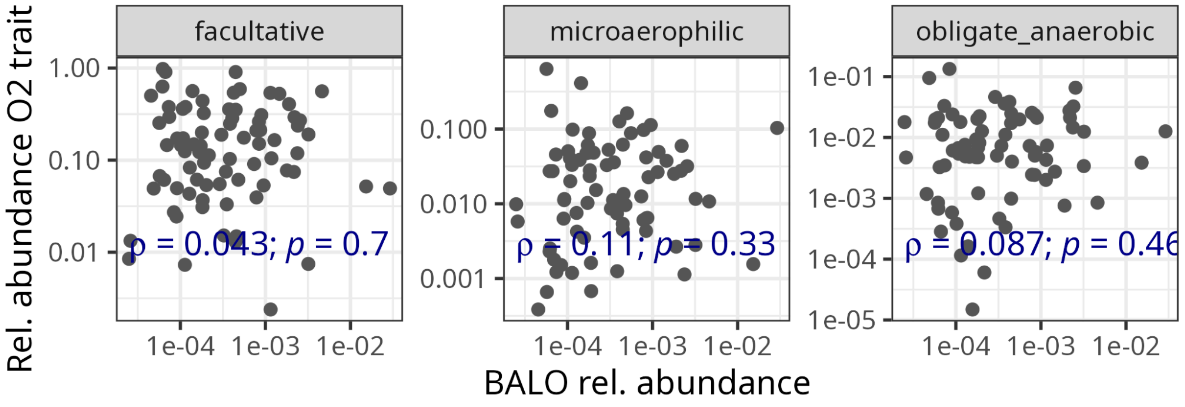


**Fig. S13: Correlation of relative abundance of oxygen preferences aggregated for each sample from species traits correlating with BALO abundance.** The comparison is between the median abundance of BALO samples for each study (n=83) of the HMC dataset. The oxygen preference per species is predicted from the 16S sequence using AmpliconTraits and summed for each sample.

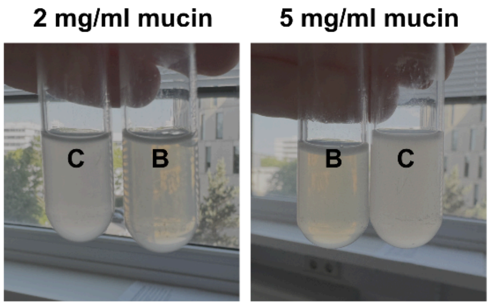


**Fig. S14: *B. bacteriovorus* HD100 can prey on *E. coli* ML35 in mucus medium.** C: control culture containing *E. coli* ML35 without the BALO. B: Co-culture with BALO *B. bacteriovorus* HD100 and *E. coli* ML35. The co-culture clearance indicates predation.

**Tab. S1: Network characteristics of networks containing samples with BALO presence versus samples without BALO presence.** Only ASVs with a mean relative abundance ≥ 0.001% were considered. Edges were filtered based on a mean correlation of at least 0.3 and a p-value > 0.05.

| | **with BALO** | **without BALO** |
|---|---|---|
| **Number of nodes** | 333 | 136 |
| **Number of edges** | 5,118 | 1,768 |
| **Avg. number of neighbors** | 32.158 | 26.758 |
| **Characteristic path length** | 2.798 | 2.905 |
| **Clustering coefficient** | 0.654 | 0.730 |
| **Network density** | 0.102 | 0.204 |
| **Network heterogeneity** | 0.880 | 0.891 |
| **Network centralization** | 0.303 | 0.351 |
| **% negative edges** | 12.16 | 1.02 |

### Tab. S2: BALO prey range according to literature

| BALO | Prey species | System | Prey is facultative | reference |
|---|---|---|---|---|
| *Bdellovibrio bacteriovorus* | *Klebsiella pneumoniae* | Rat lung | yes | Shatzkes et al. (2016) [1] |
| | *Yersinia pestis* | Mouse | yes | Russo et al. (2018) [2] |
| | *Escherichia coli* | Mouse eye, Wax-moth larvae | yes | Ottaviani et al. (2019) [3] |
| | *Pseudomonas aeruginosa* | Rabbit eye, Mouse | yes | Romanowski et al. (2023), Tajabadi et al. (2023) [4, 5] |
| | *Shigella flexneri* | Zebrafish | yes | Willis et al. (2016) [6] |
| | *Vibrio vulnificus* | Mouse | yes | Liu et al. (2023) [7] |
| | *Salmonella enterica* | Chick | yes | Atterbury et al. (2011) [8] |
| | *Pectobacterium carotovorum* | Potato | yes | Youdkes et al. (2020) [9] |
| | *Pseudomonas tolaasii* | Mushroom | obligate aerobic | Saxon et al. (2014) [10] |
| | *Pseudomonas savastanoi* pv*. glycinea* | Soybean | obligate aerobic | Scherff (1973) [11] |
| | *Vibrio cholera* | Shrimp | yes | Cao et al. (2015) [12] |
| | *Erwinia amylovora* | ? | yes | Starr and Baigent (1966) [13] |
| *Bdellovibrio* sp. | *Aeromonas hydrophila* | Fish | yes | Cao et al. (2012) [14] |
| *Bdellovibrio W* | *Rhodospirillum rubrum, Spirillum serpens* | ? | yes | Starr and Seidler (1971), Burger et al. (1968) [15, 16] |
| *Bdellovibrio krueschi* | *Ochrobactrum* | *C. elegans* | aerobic (unsure) | Maher et al. (2025) [17] |
| *Micavibrio aeruginosavorus* | *Klebsiella pneumoniae* | Rat lung | yes | Shatzkes et al. (2016) [1] |
| *Halobacteriovorax* sp. | *Vibrio* sp. | Spiny lobster | aerobic (unsure) or facultative | Ooi et al. (2021) [10] |

| | | | | |
|---|---|---|---|---|
| | *Vibrio parahaemolyticus* | Mussel | yes | Ottaviani et al. (2020) [3] |
| | *Pseudomonas* | Soil | aerobic (unsure) or facultative | Enos et al. (2018) [18] |
| | *Escherichia* | lab strain ML35 | yes | Enos et al. (2018) [18] |
| *Bacteriovorax* sp. | *Vibrio alginolyticus* | Shrimp larvae | yes | Wen et al. (2014) [19] |
| | *Vibrio parahaemolyticus* | Shrimp | yes | Kongrueng et al. (2017) [20] |
| *Pseudobdellovibrio exovorus* | Caulobacter crescentus (=Caulobacter vibrioides) | Sewage | obligate aerobe | Koval et al. (2013) [21] |
